# Dynamic functional connectivity links with treatment response of electroconvulsive therapy in major depressive disorder

**DOI:** 10.1101/2021.03.31.437958

**Authors:** Mohammad S. E. Sendi, Hossein Dini, Jing Sui, Zening Fu, Randall Espinoza, Katherine Narr, Shile Qi, Christopher C. Abbott, Sanne van Rooij, Patricio Riva-Posse, Helen S. Mayberg, Vince D. Calhoun

## Abstract

**Background:** Electroconvulsive Therapy (ECT) is one of the most effective treatments for major depressive disorder (DEP). There is recently increasing attention to evaluate ECT’s effect on resting-state functional magnetic resonance imaging (rs-fMRI). This study aims to compare rs-fMRI of DEP patients with healthy participants, investigate whether dynamic functional network connectivity network (dFNC) estimated from rs-fMRI predicts the ECT outcome, and explore the effect of ECT on brain network states.

**Method:** Resting-state fMRI data were collected from 119 patients with depression or DEP (76 females), and 61 Healthy (HC) participants (34 females) with an age mean of 52.25 (N=180) years old. The pre-ECT and post-ECT Hamilton Depression Rating Scale (HDRS) were 25.59±6.14 and 11.48±9.07, respectively. Twenty-four independent components from default mode (DMN) and cognitive control network (CCN) were extracted using group-independent component analysis from pre-ECT and post-ECT rs-fMRI. Then, the sliding window approach was used to estimate the pre-and post-ECT dFNC of each participant. Next, k-means clustering was separately applied to pre-ECT dFNC and post-ECT dFNC to assess three distinct states from each participant. We calculated the amount of time each individual spends in each state, called occupancy rate or OCR. Next, we compared OCR values between HC and DEP participants. We also calculated the partial correlation between pre-ECT OCRs and HDRS change while controlling for age, gender, number of treatment, and site. Finally, we evaluated the effectiveness of ECT by comparing pre-and post-ECT OCR of DEP and HC participants.

**Results:** The main findings include: 1) DEP patients had significantly lower OCR values than the HC group in a state, where connectivity between CCN and DMN was relatively higher than other states (corrected p= 0.015), 2) Pre-ECT OCR of state, with more negative connectivity between CCN and DMN components, predicted the HDRS changes (R=0.23 corrected p=0.03). This means that those DEP patients who spend less time in this state showed more HDRS change, and 3) The post-ECT OCR analysis suggested that ECT increased the amount of time DEP patients spend in state 2 (corrected p=0.03). Finally, we found ECT increases the total traveled distance in DEP.

**Conclusion:** Our finding suggests that dFNC features, estimated from CCN and DMN, show promise as a predictive biomarker of the ECT outcome of DEP patients. Also, this study identified a possible underlying mechanism associated with the ECT effect in DEP patients.

## 1 Introduction

Major depressive disorder (MDD) is a debilitating brain disorder (Tsuchiyagaito et al., 2021), which is characterized by impaired cognitive functioning, somatic abnormalities such as inattention and inability to focus, as well as emotional troubles (Liu et al., 2021; Luo et al., 2021). Based on the global burden of disease report, depression is the third rank cause of making people disabled, and it has been estimated to be the first rank cause of burden before 2030 (Organization, 2008, Ebneabbasi et al., 2021). There are effective treatments accessible such as psychotherapy and alleviating medications, but it has been declared that 30 percent of patients suffering from MDD do not respond to these treatments (Rush et al., 2006). Therefore, there is an essential need for advanced therapies such as deep brain stimulation (DBS), transcranial magnetic stimulation (TMS), and electroconvulsive therapy (ECT) which are being broadly used as an alternative to alleviate MDD symptoms (Settell et al., 2017; Mo et al., 2020; Williams et al., 2021).

Among all mentioned therapies, ECT can be considered one of the most effective treatments for pharmacological resistant MDD (Enneking et al., 2020) due to its faster action and higher remission rate than typical medicine-based treatments (Leaver et al., 2020). One hundred thousand annual ECT treatments in the U.S revealed that this treatment’s success rate is around 75% (Abbott et al., 2013). It has been shown that most depressive episodes disappeared after 3-4 weeks of ECT series (Abbott et al., 2013). Yet, underlying neural and cognitive mechanisms behind this improvement caused by ECT are still unclear. Moreover, the question of whether we can predict the effectiveness of ECT before applying it remained open.

In recent years, functional network connectivity (FNC) data obtained from resting-state functional magnetic resonance imaging (rs-fMRI) time series has demonstrated highly informative about the underlying brain connectivity patterns in mental disorders such as MDD (Mulders et al., 2015; Yan et al., 2019; Liu et al., 2020; Luo et al., 2021). Recently, studies have shown that ECT resets and stimulates the formation of the brain regions/networks connectivity (Wang et al., 2018a, 2018b; Bai et al., 2020). An investigation of the whole-brain FNC of the patients with depression showed a reduction in the left dorsal lateral prefrontal cortex connectivity corresponded to ECT therapeutic course (Perrin et al., 2012). Several recent studies reported functional and structural connectivity changes occurred in the amygdala and anterior cingulate cortex (ACC) after ECT (Wang et al., 2017; Takamiya et al., 2018; Gryglewski et al., 2019; Qiu et al., 2019; Sartorius et al., 2019). Another study declared that the cognitive control network (CCN) and the default mode network (DMN) play a vital role as the most effective brain networks in regulating brain connections after ECT (Menon, 2015). Moreover, increased intra-network connectivity in CCN (Wang et al., 2018a, 2018b), and decreased dorsolateral prefrontal cortex global functional connectivity(DLPFC) as a part of CCN (Perrin et al., 2012), are reported to be occurred after applying ECT. Other rs-fMRI studies reported significant changes in functional connectivity of DMN by applying ECT (Mulders et al., 2016; Wei et al., 2018; Bai et al., 2019).

In the aforementioned studies, FNC estimated at CCN and DMN is often assumed to be static over time. However, this assumption runs contrary to the dynamic nature of brain FNC, and dynamic FNC (dFNC) has been recently introduced to overcome this limitation (Allen et al., 2014; Elaheh Zendehrouh et al., 2020; Sendi et al., 2021c). In recent years, a few studies have looked into dFNC in MDD and discovered that MDD affects both the strength and the temporal properties of FNC. As a result, we hypothesized that looking at the effect of ECT on dFNC might reveal how and to what extent ECT affects DMN and CCN.

In this study, we used rs-fMRI data from 119 patients with depression (DEP) who experienced a series of ECT and 61 healthy controls (HC) to find the neural mechanisms behind the improvement after ECT and predict the effectiveness of ECT before applying it. To this aim, we used group independent component analysis (ICA) and extracted independent components from DMN and CCN and estimated dFNC in these two networks by applying sliding window approach and clustered dFNC into a few brain states using k-means clustering. Finally, we compared the occupancy rate (OCR) estimated from state vector, an output of k-means clustering, between HC and DEP in both pre-and post-ECT. Moreover, correlating OCR with clinical data, we could significantly predict the effectiveness of ECT by using just pre-ECT rs-fMRI data.

## 2 Materials and Methods

### 2.1 Participants and clinical outcome

This study used neuroimaging, clinical, and demographic information of 119 patients (76 females) diagnosed with depression (called DEP hereafter) and 61 healthy (HC) participants (34 females) from either University of New Mexico (UNM) or the University of California Los Angeles (UCLA). Exclusion criteria were as follows: 1) Having any neurodegenerative and neurological disorders such as Alzheimer’s disease or psychiatric conditions such as schizophrenia.2) Having alcohol or drug addiction, pregnancy, and potential dangers under magnetic resonance imaging (MRI) such as using a pacemaker.

Hamilton Depression Rating Scale-17 items (HDRS) were used to assess the patient group’s symptom severity before and after the ECT (Heijnen et al., 2010). Initial and final assessments were given to participants before ECT starts and within a week of completing ECT series, respectively. Some participants from UNM were investigated to have psychotropic medications initially, but further investigations on UNM and all UCLA participants they stopped revealed psychotropic medications before ECT outset. Demographical information, and clinical measurements can be seen in Table1. Finally, All the participants signed the consent form, and this study has been approved by the institutional review boards at UNM and UCLA.

**Table 1.**
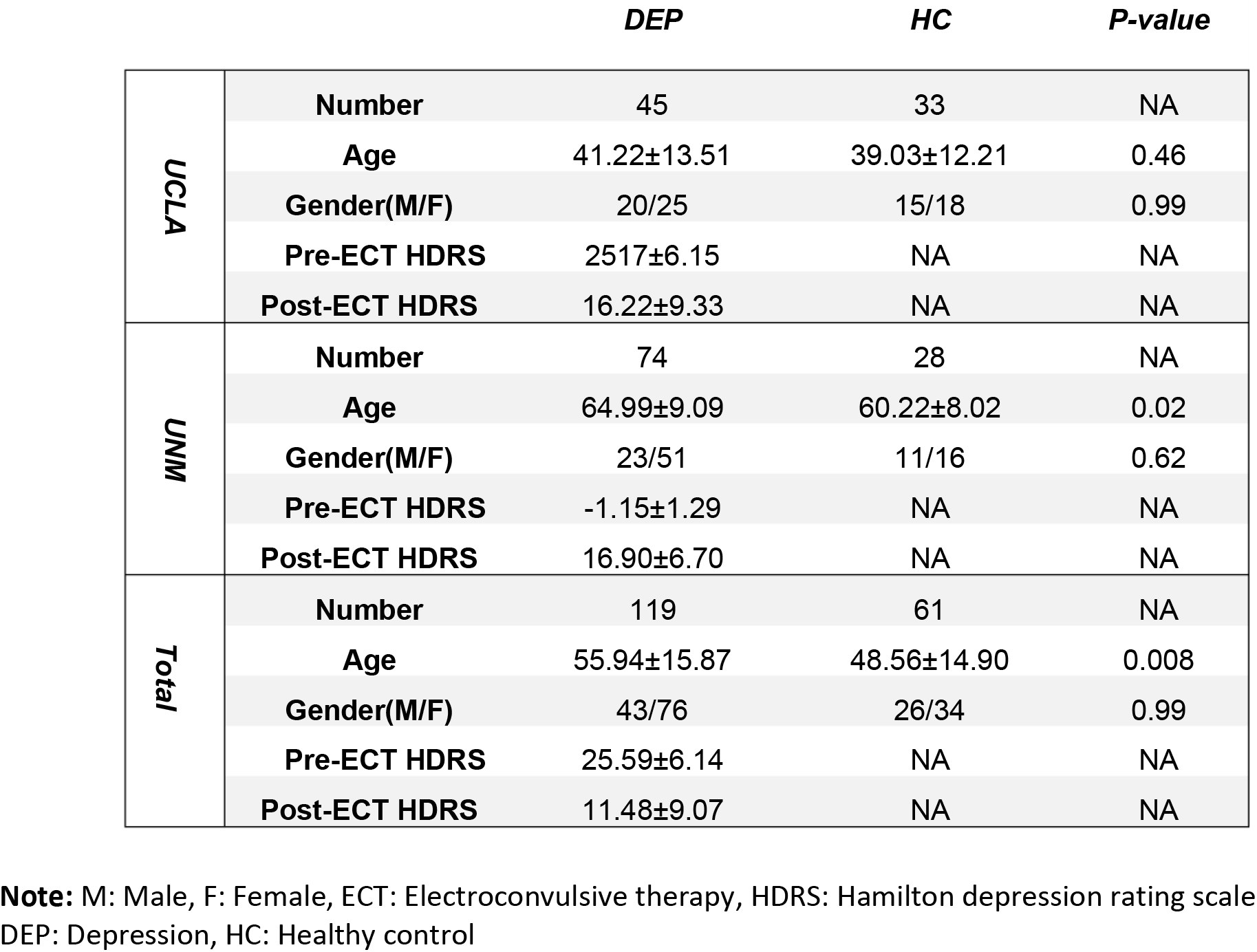
Demographic and clinical details of participants for each site.

### 2.2 ECT procedure

In the UNM site, Thymatron System IV (Somatics, Lake Bluff, IL, USA) was used, and the ECT procedure was initiated with a right unilateral d’Elia (ultra-brief pulse width of 0.3 ms, stimulus dosage at 6 × threshold) placement of electrodes. If the participant did not respond to ECT, the treatment continued with bitemporal (brief pulse width of 1 ms, stimulus dosage at 2 × threshold) electrode placement. In UCLA, a Mecta 5000Q (MECTA Corp., Tualatin, OR, USA), the exact electrode placement, and similar stimulus dosages are used in UNM. Treatments were applied three times a week, and until obtaining a stable clinical response or psychiatrist decision to stop treatment in the context of nonresponse. ECT implementation procedure followed the clinical standards announced by the APA ECT Task Force Report and was not manipulated for the goal of this study. During the treatment process, patients were oxygenated and received adequate induction (methohexital or etomidate) and relaxation (succinylcholine). Clinical measures such as blood pressure were monitored during the treatment.

### 2.3 fMRI data acquisition

At the UNM site, a 3-Tesla Siemens Trio scanner (Siemens Healthcare, Malvern, PA, USA) was used to collect MRI data. Parameters of the whole-brain gradient-echo echo-planar imaging sequence are as follows: echo time (TE) = 29 milliseconds (ms), repetition time (TR) = 2 s (s), voxel size = 3.75 × 3.75 × 4.55 mm, flip angle (FA) 75°, and 154 volumes. At UCLA, a 3-Tesla MAGNETOM Allegra MRI scanner (Siemens, Erlangen, Germany) was used to collect MRI data. Parameters of functional images are as follows: TE = 30 ms, TR = 5 s, voxel size= 3.4 × 3.4 × 5 mm, FA = 70°, and 180 volumes. The duration of resting-state scans was a minimum of 5 minutes and 16 seconds, and participants were guided to passively keep their concentration to the fixation cross during the scan.

### 2.4 Data preprocessing

Here are the standard preprocessing steps for fMRI data using statistical parametric mapping (SPM12, https://www.fl.ion.ucl.ac.uk/spm/): 1) to address longitudinal relaxation effects, five initial fMRI scans were removed. 2) time differences in slice acquisition were corrected. 3) motion was corrected using SPM. 4) imaging data was spatially normalized based on echo-planar imaging (EPI) template in standard Montreal Neurological Institute (MNI) space and was resampled to 3×3×3 mm^3^. 5) a 6-mm full-width half-maximum (FWHM) Gaussian kernel for spatial smoothing is applied to the data.

The Neuromark fully automated group ICA pipeline using GIFT (http://trendscenter.org/software/gift) is implemented to extract reliable CCN and DMN independent components (ICs). In this method, previously derived components maps are used as priors for spatially constrained ICA; considering group-level spatial maps from two large-sample HC datasets, replicable components were identified and used for the spatial priors (Du et al., 2020). Prior to the dynamic functional connectivity method, we implement some other de-noising and artifact rejection steps as follows: 1) linear, quadratic, and cubic de-trending. 2) use of 6 realignment parameters and their temporal derivatives for multiple regression 3) outlier removal and band-pass filtering (from 0.01 to 0.15 Hz). Identified components in cognitive control network (CCN) and default mode network (DMN) can be seen in Table2.

**Table 2.** Component Labels.

|  |  | Component name | Peak coordinate (mm) |  |  |
| --- | --- | --- | --- | --- | --- |
| 1 | CCN | Inferior parietal lobule ([IPL], 68) | 45.5 | -61.5 | 43.5 |
| 2 |  | Insula (33) | -30.5 | 22.5 | -3.5 |
| 3 |  | Superior medial frontal gyrus ([SMFG], 43) | -0.5 | 50.5 | 29.5 |
| 4 |  | Inferior frontal gyrus ([IFG], 70) | -48.5 | 34.5 | -0.5 |
| 5 |  | Right inferior frontal gyrus ([R IFG], 61) | 53.5 | 22.5 | 13.5 |
| 6 |  | Middle frontal gyrus ([MiFG], 55) | -41.5 | 19.5 | 26.5 |
| 7 |  | Inferior parietal lobule ([IPL], 63) | -53.5 | -49.5 | 43.5 |
| 8 |  | Left inferior parietal lobue ([R IPL], 79) | 44.5 | -34.5 | 46.5 |
| 9 |  | Supplementary motor area ([SMA], 84) | -6.5 | 13.5 | 64.5 |
| 10 |  | Superior frontal gyrus ([SFG], 96) | -24.5 | 26.5 | 49.5 |
| 11 |  | Middle frontal gyrus ([MiFG], 88) | 30.5 | 41.5 | 28.5 |
| 12 |  | Hippocampus ([HiPP], 48) | 23.5 | -9.5 | -16.5 |
| 13 |  | Left inferior parietal lobue ([L IPL], 81) | 45.5 | -61.5 | 43.5 |
| 14 |  | Middle cingulate cortex ([MCC], 37) | -15.5 | 20.5 | 37.5 |
| 15 |  | Inferior frontal gyrus ([IFG], 67) | 39.5 | 44.5 | -0.5 |
| 16 |  | Middle frontal gyrus ([MiFG], 38) | -26.5 | 47.5 | 5.5 |
| 17 |  | Hippocampus ([HiPP], 83) | -24.5 | -36.5 | 1.5 |
| 18 | DMN | Precuneus (32) | -8.5 | -66.5 | 35.5 |
| 19 |  | Precuneus (40) | -12.5 | -54.5 | 14.5 |
| 20 |  | Anterior cingulate cortex ([ACC], 23) | -2.5 | 35.5 | 2.5 |
| 21 |  | Posterior cingulate cortex ([PCC], 71) | -5.5 | -28.5 | 26.5 |
| 22 |  | Anterior cingulate cortex ([ACC], 17) | -9.5 | 46.5 | -10.5 |
| 23 |  | Precuneus (51) | -0.5 | -48.5 | 49.5 |
| 24 |  | Posterior cingulate cortex ([PCC], 94) | -2.5 | 54.5 | 31.5 |

### 2.5 Functional network connectivity

A sliding window which is a convolution of a rectangle (window size = 20 TRs = 40 s) with a Gaussian (σ = 3s), applied to the data to calculate DEP. This method was used to localize the dataset per time point, and the procedure can be seen in Fig1 Step1. Since the neuroimaging data have different temporal resolutions (TR = 2 s for UNM and 5 s for UCLA), the individual data with the low temporal resolution was interpolated based on the high temporal resolution data to construct new TCs. Such an interpolation strategy has been successfully introduced in previous dFNC studies, showing reliable performance for capturing FNC dynamics in datasets with different temporal resolutions (de Lacy et al., 2017; Fu et al., 2019). Next, we used the Pearson correlation method to calculate dFNC between 24 sub-nodes of DMN and CCN. Then, we obtained 276 connectivity features (Fig1 Step1). Calculated dFNC for each window was concatenated for each individual as a form of C × C × T array (where C is the number of ICs and equals 276, and T represents total windows and equals 610). Finally, all arrays for all participants were concatenated to show brain connectivity changes between ICs as a function of time (Fig1 step2) (Allen et al., 2014; Sendi et al., 2021b, 2021a).

**Fig. 1.**
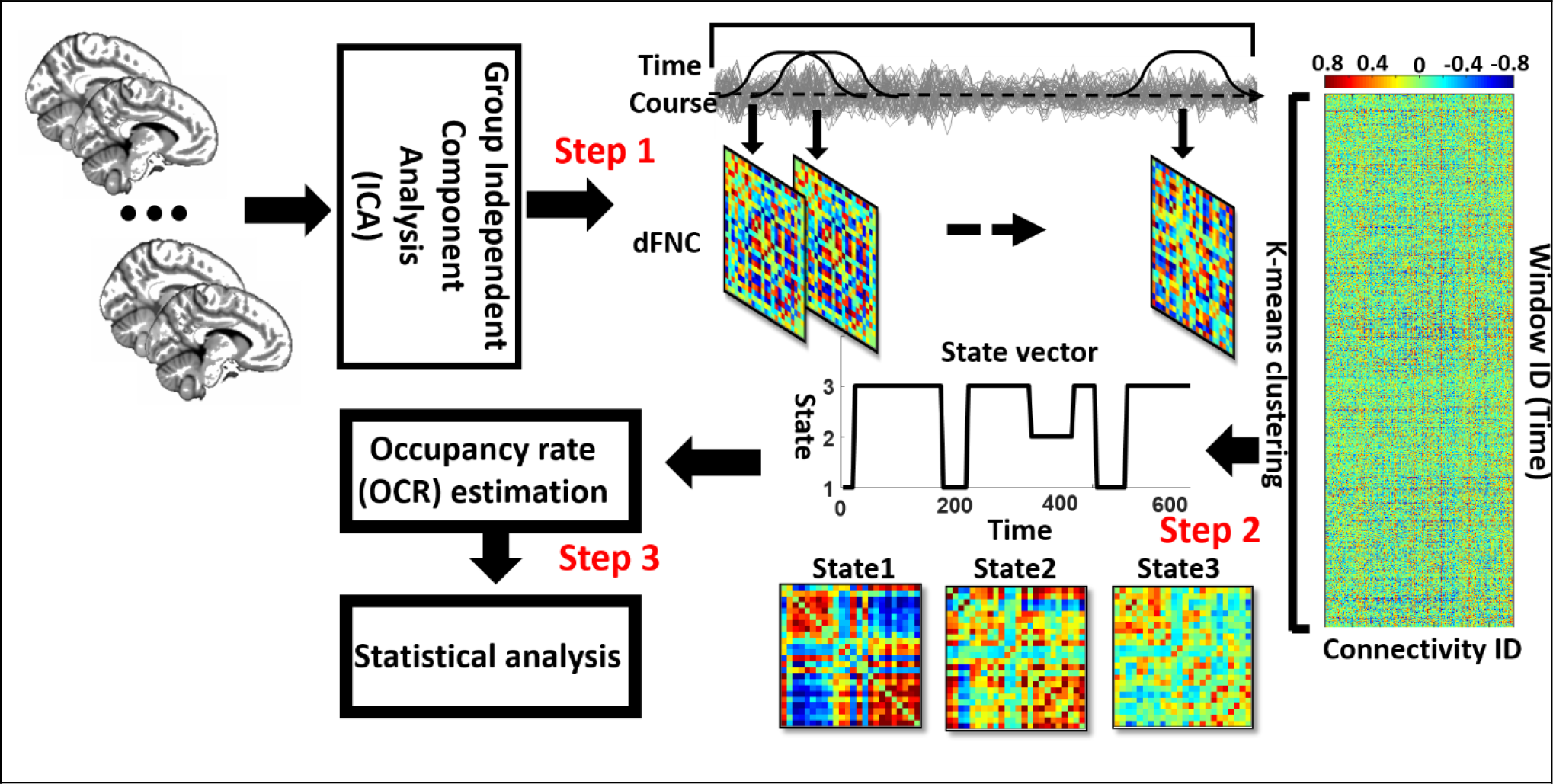
Analytic pipeline: The time-course signal of 24 components of the CCN/DMN networks has been identified using group independent component analysis (ICA). In step2, a taper sliding window was used to segment the time-course signals and then calculated the functional network connectivity (FNC). After vectorizing the FNC matrixes, we have concatenated them, and then a k-means clustering, k=3, was used to group FNCs to three distinct states (Step2). Elbow criteria were used to find the optimal k. In addition, the correlation distance metric is used in this clustering. Then, based on the state vector of each participant, the occupancy rate or OCR features, in total 3 features, were calculated from the state vector of each participant. Then we compared the OCR among groups. By using ttest. We adjusted all p values by the Benjamini-Hochberg false discovery rate (FDR) correction in each analysis (Step3).

### 2.6 Clustering and dFNC latent features

We implemented a K-means clustering method on the previous step’s output. First, we concatenated dFNCs of all participants, then we used a k-means clustering method to put them into a few clusters or states (Allen et al., 2014; Sendi et al., 2021c). We used the elbow criterion to calculate the optimum number of clusters (optimum k in the k-means method), a clustering analysis standard (Sendi et al., 2021c). This method defines the optimization equation as the distance of within-cluster and between clusters as a ratio and tries to minimize this ratio. We found the optimal number of clusters is 3, searching from k=2 to 9. We used the L1 norm as our distance metric with 1000 iterations. This process yielded 3 distinct states for the group of participants and the state vector for each individual. State vector shows the state of brain at any given time. Subsequently, based on the state vector, we calculated each subject’s time interval (the number of time windows that each participant was in a specific state), and we call this feature the OCR of each state (Fig1 Step3). Thus, considering three states, we have three OCR for each individual. Finally, we calculated the traveled distance for each participant using Euclidean distance. To determine the distance traveled, we calculated the distance between each time window of dFNC matrix and then summed up all possible window pairs’ distance. So, we have one traveled distance for each individual, and it is a state-independent metric.

### 2.7 Statistical analysis

The OCR feature and traveled distance between DEP and HC group is compared using two samples t-test. Moreover, to see whether OCR can predict HDRS scores, partial correlation accounting for age, gender, number of treatments, and scanning site is used. All p values were adjusted by the Benjamini-Hochberg method for false discovery rate or FDR (Yoav Benjamini; Yosef Hochberg, 1995).

## 3 Results

This section discusses the results obtained from dFNC analysis and the comparison between DEP and HC groups. It consists of dFNC states resulting from clustering analysis, the correlation of OCR with HDRS scores, comparison between DEP and HC in OCR both in pre-ECT and post-ECT, and the traveled distance between two groups.

### 3.1 Clinical results

The clinical and demographical information of participants is provided in Table 1 separately for DEP and HC groups. Concerning the HDRS scores, the DEP group has an average of 25.59 and a standard deviation of 6.14 in pre-ECT, an average of 11.48, and a standard deviation of 9.07 in post-ECT. Using a two-sample t-test, we found a significant difference (p<0.00) between HDRS values of pre- and post-ECT values. In other words, the clinical scores showed that after implementation of ECT, the HDRS scores had significantly decreased.

### 3.2 Dynamic functional network connectivity states for pre-ECT and post-ECT

We found three separate clusters (states) applying the k-means clustering method on dFNC of all individuals (both DEP and HC group). It worth mentioning that we applied the clustering method to pre-ECT and post-ECT dFNC data separately. Fig2.A and Fig2.B show these three distinct states for pre-ECT and post-ECT, respectively. We found pre-ECT and post-ECT rs-fMRI generate similar brain states. To assess this similarity across corresponding states, we used Fisher correlation coefficients (state1: R=98.47%, state2: R=87.5%, state3: R=97.67%). Additionally, we calculated the average of dFNC values of CCN, DMN, and CCN/DMN (i.e., the connectivity between DMN and CCN) as shown in Table 3.

**Fig2.**
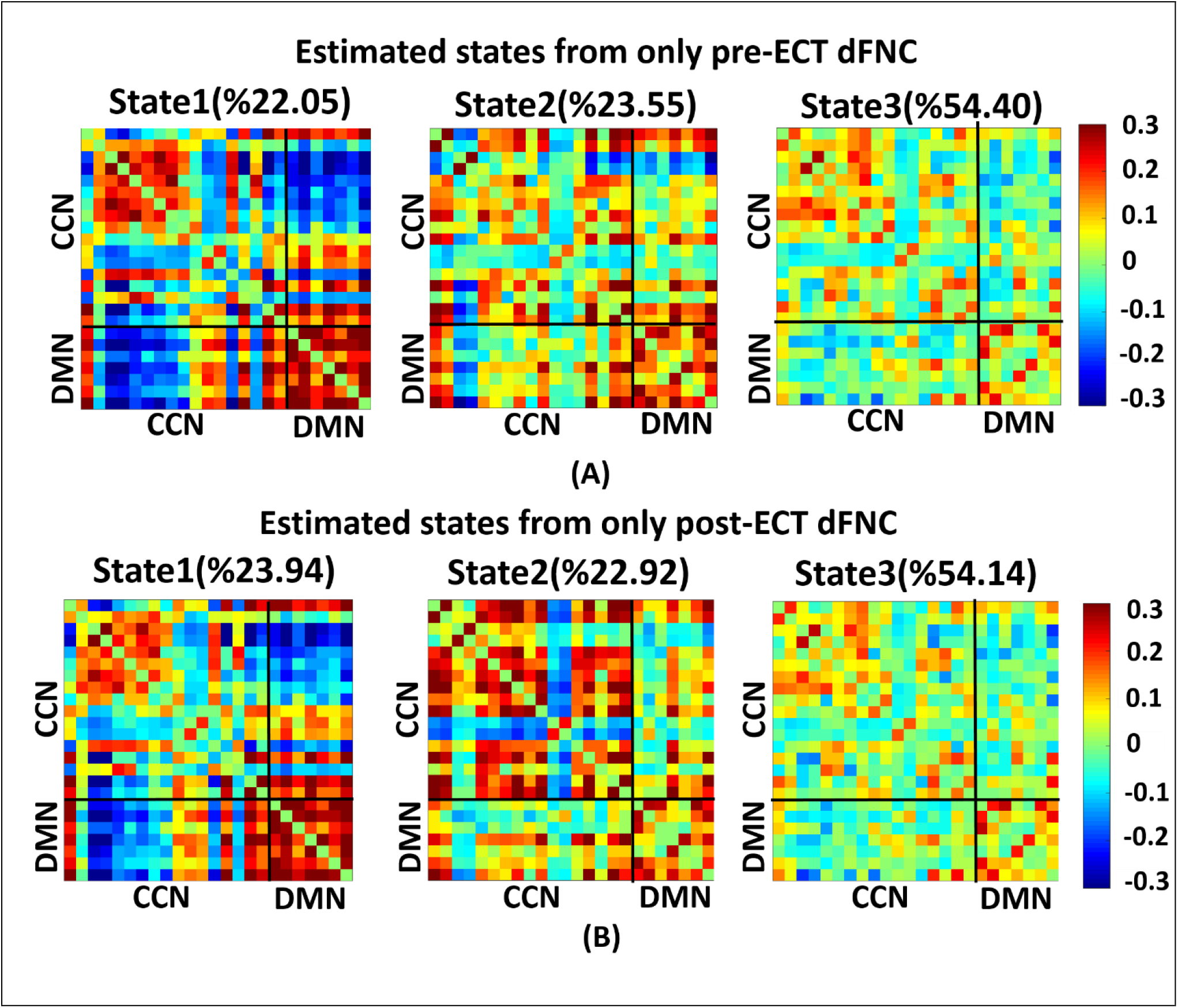
Dynamic functional connectivities in three identified states using clustering method. Each state is consisted of a 24×24 matrix where the positive connectivities are shown by hot color and negative connectivities are shown with cold colors. The values in parentheses show the overall percentage of time participants spent in each specific state. **A)** States resulted from clustering analysis on pre-ECT dFNC. **B)** States resulted from clustering analysis on post-ECT dFNC.

**Table 3.**
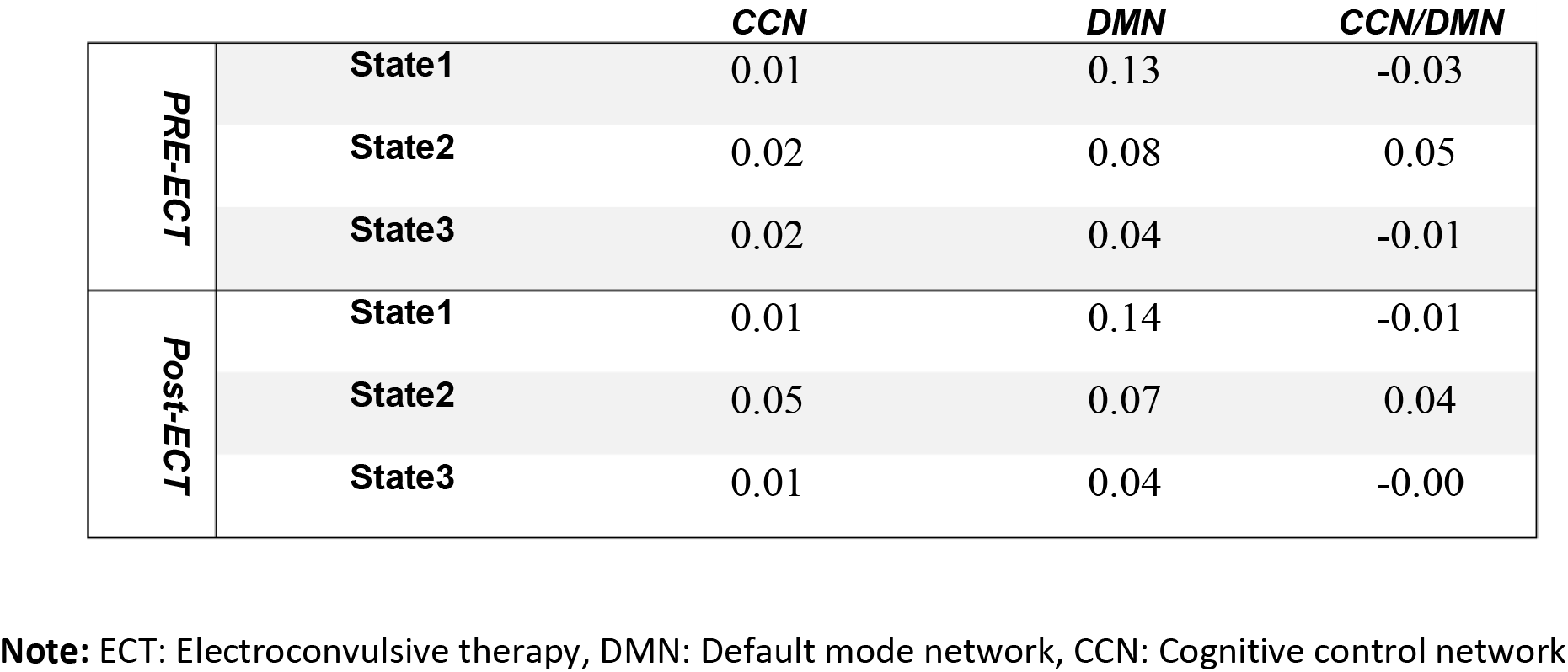
The state-specific dFNC average.

We found that both state 2 and state 3 have higher within-CCN connectivity than state 1 in pre-ECT. While state 1 had more increased within-DMN connectivity than the other two states in both pre-ECT and post-ECT. Only state 2 showed positive functional connectivity between DMN and CCN in both pre-ECT and post-ECT data.

### 3.3 Comparison of OCR and travelled distance between HC and DEP in pre-ECT and post-ECT

Fig3.A and Fig.3B show the OCR values of HC and DEP groups in different states for pre-ECT and post-ECT, respectively. Only OCR of state 2, with relatively higher CCN/DMN functional connectivity, shows a significant difference between DEP and HC. We found HC spend more time in state2 before applying ECT (FDR corrected p=0.015), while the pattern in reversed after ECT and patients with depression spend more time in state 2 (FDR corrected p=0.03). Moreover, we compared the traveled distance between DEP and HC groups in pre-ECT and post-ECT (Fig 3.C). The results showed that in pre-ECT, HC group traveled significantly more distance compared to DEP group (p=0.04). In post-ECT, again, the HC group’s traveled distance is higher than DEP group, but this difference is not significant.

**Fig3:**
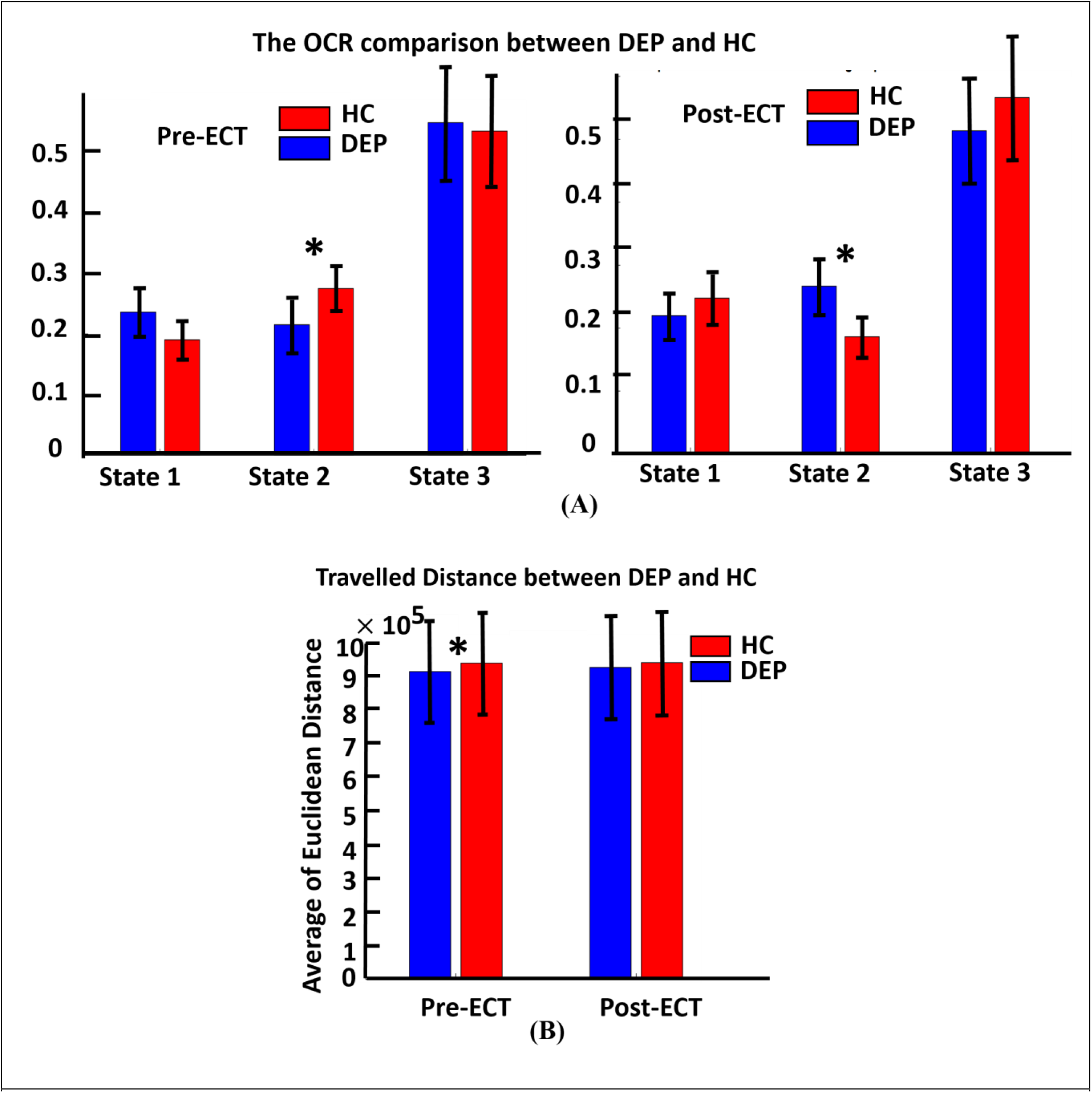
**(A)** The OCR comparison between DEP and HC in three distinct states of pre-ECT (left image) and post-ECT (right image). Red bars indicating average OCR for healthy group in each state and blue bars are for the OCR of DEP group in each state. In the left image, OCR features are extracted from dFNC of just pre-ECT (significant difference in state2, corrected p=0.015). In the right image, OCR features extracted from dFNC of just post-ECT(significant difference in state2, corrected p=0.03). In state2, ECT had significantly changed the OCR value of HC and DEP before applying ECT (HC>DEP) compared to after applying ECT (HC<DEP). **B)** shows the travelled distance between DEP and HC group in pre-ECT and post-ECT. In pre-ECT, the travelled distance of HC group is significantly higher than DEP group (p=0.04). After applying ECT, HC group has higher travelled distance than DEP group but this difference is not significant.

### 3.4 The link between Pre-ECT OCR and the effectiveness of ECT

To predict whether applying ECT would be effective, we correlated the calculated OCR of 119 patients with their associated HDRS change (post_HDRS-pre_HDRS) by controlling the age, gender, scanning site the number of treatments. As shown in Fig.4, only the OCR of state 1 is a significant predictor (R=0.22, FDR corrected p=0.03. In more detail, we found those patients who spend more time in state 1, with relatively lower CCN/DMN functional connectivity, showed less reduction in their HDRS. We did not find a significant link between the pre-ECT traveled distance and the HDRS change.

**Fig4:**
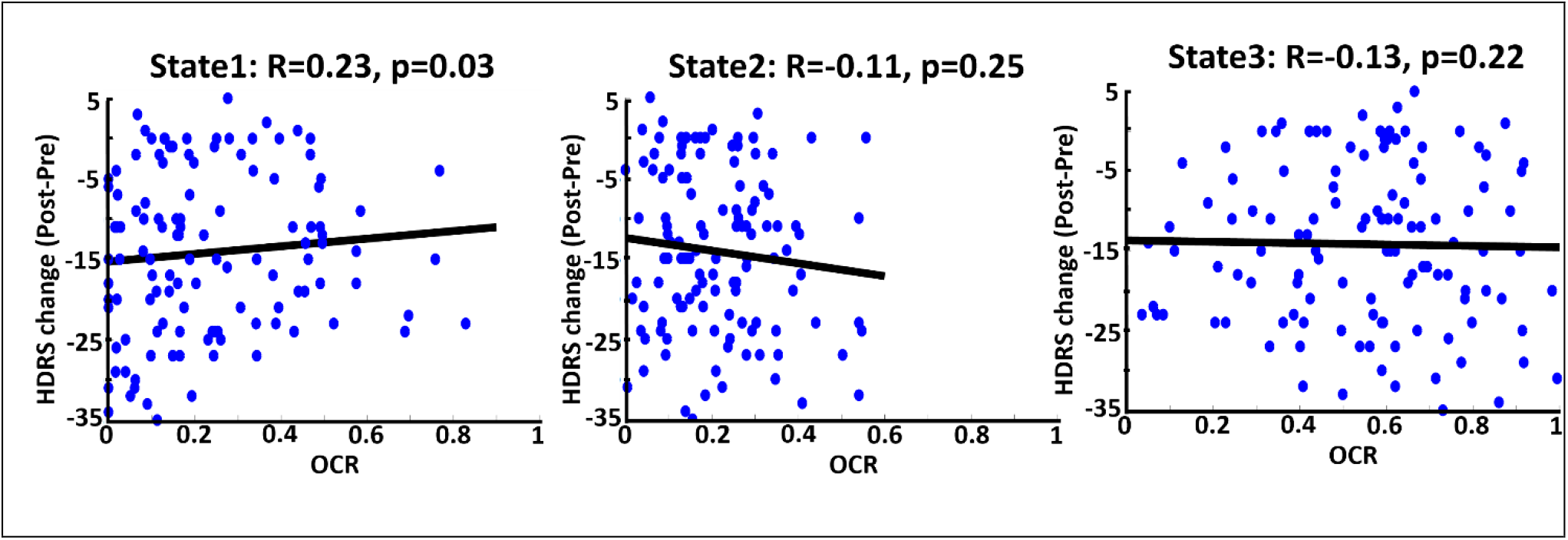
Correlation between OCR values and reported HDRS change (Pre-Post) in three states. Blue dots are referred to 119 DEP individuals. The bold black line is the fitted curve. R indicates the fitted line slope in each state. As it is shown, state 1 (state with the lowest CCN/DMN connectivity) significantly predicts the HDRS change based on OCR values, less OCR value corresponds to more HDRS change.

## 4 Discussion

In this study, we used rs-fMRI of 119 participants suffering from depression and experiencing series of ECT treatment and 61 healthy individuals to predict the effectiveness of ECT and investigate brain dynamics in pre-ECT and post-ECT by comparing DEP and HC group. To this aim, we used dFNC features extracted from CCN and DMN and clustered them into three different states. In this study, using a data-driven method of analyzing dFNC in CCN and DMN, we demonstrated that these brain networks are thoroughly dynamic in both pre-and post-ECT states. This finding agrees with previous studies on MDD that have provided evidence of dynamism in CCN and DMN (Repple et al., 2020; Yang et al., 2020; Chen et al., 2021; Sendi et al., 2021a).

We compared the differences between DEPs and HC groups considering this temporal dynamic activity of the brain. To this aim, we calculated the OCR feature, which shows the proportional amount of time participants spent in each state. Comparing the OCR of DEP with HC group in pre-ECT, we found that the OCR of HC group is significantly higher than the DEP group in state 2. This means that HC group tends to spend more time in state 2 than DEPs. The main characteristic of pre-ECT state 2 is having higher CCN/DMN connectivity than other pre-ECT states.

The difference between MDD and HC group with a specific focus on DMN and CCN has been reported in previous literature. For example, one study reported an increase in within-DMN connectivity for MDD (Posner et al., 2016), while another study reported a decrease in this network’s connectivity (Li et al., 2020; Wang et al., 2020). Additionally, recent studies using rs-fMRI suggest an association between depression and abnormal functional connectivity in CCN network (Schlösser et al., 2008; Vasic et al., 2009; Sheline et al., 2010; Veer et al., 2010; Alexopoulos et al., 2013; Clasen et al., 2014). Other studies focusing on the CCN network reported attenuated connectivity of that network in remitted MDDs (Stange et al., 2017; Jiao et al., 2020). Finally, a study tried to predict the antidepressant response in MDDs focusing on within-CCN and within-DMN networks and reported low and high resting functional connectivity within-CCN and within-DMN, respectively (Alexopoulos et al., 2012). While previous studies focused only on within-DMN and within-CCN functional connectivity, the current study might provide new evidence about the role of CCN/ DMN connectivity in depression.

Furthermore, we investigated the effect of ECT on evaluating the temporal dynamic activity of the brain after implementing ECT in post-ECT. We found the OCR of DEPs is significantly higher than HC participants. This means that after ECT, patients with depression spent more time in state 2 than HC group. Similar to pre-ECT state2, post-ECT state2 shows the highest CCN/DMN connectivity than other states. That might provide new insight into the effect of ECT on the CCN and DMN by regulating the temporal dynamics of these brain networks. Moreover, post-ECT state2 has relatively higher within-CCN connectivity. This contrasts to previous studies that reported reduced within-CCN connectivity associated with the antidepressant state in a relatively small dataset (Alexopoulos et al., 2012); we found increased within-CCN connectivity after ECT.

Evaluating the effectiveness of ECT, i.e., identifying patients as a potential remitter or non-remitter before implementing ECT, would be valuable from the clinical perspective (Van Waarde et al., 2015). On the one hand, many studies have correlated baseline clinical characteristics with MDD status outcome (Perlman et al., 2019; Kennis et al., 2020). Such analyses are based on group-level analysis rather than individual patient-level aspects (Ozomaro et al., 2013). There is a need for new metrics to predict ECT outcomes. On the other hand, selecting a feature among many MRI metrics is difficult because they focus on non-overlapping aspects of brain function (Leaver et al., 2018). Although some metrics are based on functional connectivity of fMRI data, there are focused on static brain region connections (Leaver et al., 2018). Therefore, the use of metrics based on dFNC and using the correlation analysis of such metrics with behavioral and clinical data could help predict ECT outcome. In this study, we were able to predict the effectiveness of ECT before applying it. To this aim, we correlated the HDRS change with just pre-ECT OCR of DEPs and found that brain dynamics in state 1 is the predictor (Fig4). We found a significant correlation between OCR and HDRS change with a positive slope, which means that the more OCR in pre-ECT state 1, with relatively less CCN/DMN, equals fewer HDRS changes. Therefore, DEPs who spent more time in pre-ECT state1 are less likely to be treated by ECT. Interestingly, the main characteristic of state 1 is that this state has the lowest connectivity between CCN/DMN relative to state 2 and state 3 (Fig2-A and Table3). Also, the results of the effect of ECT showed that ECT had increased the amount of time that DEPs are spending in post-ECT state 2, with higher CCN/DMN connectivity. As such, we can conclude that the results of prediction are in line with the result of the effect of ECT, since spending time in the state with the minimum correlation between CCN andDMN is not good, and ECT had increased the amount of time DEPs spending in the state where the CCN/DMN correlation is maximum.

Finally, we extracted total travelled distance metric from pre- and post-ECT dFNC using Euclidean distance. This metric shows the dynamic of brain since it measures the distance travelled between each window’s dFNC. As shown in Fig3-B, in pre-ECT, the travelled distance of DEPs is significantly lower than HC group, but after ECT this difference is not significant. In other words, ECT decreases the traveled distance difference between HC and DEP. Therefore, we can conclude that ECT makes the brain activity of DEPs more dynamic.

### 4.1 Limitations

There are a few limitations in the current study. First, we did not directly measure whether participants were awake or closed their eyes during the scanning. Concerning this issue, we used the questionnaire and self-reports provided by participants after the scanning finished. Based on the literature, this issue might affect our results (Agcaoglu et al., 2019). To address this, it is possible to extend the dynamic functional connectivity approach to assess when the participant’s eyes were closed, or they are exhibiting aspects of drowsiness (Allen et al., 2018). dFNC approaches have already shown promise in predicting measures of drowsiness or sleep (Damaraju et al., 2020). Moreover, our data were collected in two different sites, and the imaging protocol and the number of treatments were different in these two sites. Addressing this issue, we tried to consider these differences by including site and number of treatments as a covariate in our analysis to control their effect on the results. Finally, despite the fact that HDRS is generally utilized in scaling the depression symptom severity, this score relies upon the interviewer’s skill and knowledge (Sharp, 2015). Since the data in this study comes from two separate sites, each with its own set of raters. This may cause HDRS values to vary and be inaccurate across sites.

### 4.2 Conclusion

This study evaluated dynamic functional network connectivity of DMN and CCN using rs-fMRI data of DEP patients experiencing ECT treatment. Focusing on CCN and DMN networks and clustering the brain dFNC to three different states, we found brain activity in these networks is highly dynamic. Comparing the OCR feature extracted from dFNC of these two networks between DEP and HC groups, we found that HC group prefers to spend more time in a state where the connectivity between CCN and DMN is the maximum. Moreover, we found that ECT causes an increase in the amount of time DEP patients spend in the state in which the of CCN/DMN functional connectivity is maximum. In addition, we could significantly predict the effectiveness of the ECT using just pre-ECT brain activity. We found that the more time participants spend in the state in which the correlation of CCN/DMN is minimum, the less HDRS change they have, and the less effectiveness of ECT they would experience. Finally, we found that the distance that DEP patients travel before ECT is significantly lower than the distance they travel after ECT compared to HC group. While this difference was not significant after ECT. This suggests an increase in brain dynamics after implementing ECT. In brief, this study provided a focus on functional connectivity dynamics of CCN and DMN network of DEP patients and introduced CCN/DMN connectivity as a biomarker by which we can predict the effectiveness of ECT.

## 6 Author Contributions

Mohammad S. E. Sendi and Hossein Dini developed the study, conducted data analysis, interpreted the results, and wrote the original manuscript draft. Zening Fu, Shile Qi preprocessed the data. Randall Espinoza, Katherine Narr collected the data. Christopher C. Abbot collected the data and provided a critical review of the initial draft. Sanne van Rooij, Patricio Riva-Posse, and Helen S. Mayberg provided a critical review of the initial draft. Vince D. Calhoun developed the study, interpreted the results, edited the original draft, and provided critical review to the initial draft. All authors approved the final manuscript.

## 7 Data and code Availability Statement

The code used for preprocessing and dFNC calculation are available at https://trendscenter.org/software/

## 8 Acknowledgments

We thank those who helped collect this valuable data.

## Reference

Abbott, C. C., Lemke, N. T., Gopal, S., Thoma, R. J., Bustillo, J., Calhoun, V. D., et al. (2013). Electroconvulsive therapy response in major depressive disorder: a pilot functional network connectivity resting state fMRI investigation. 4, 1–9. doi:10.3389/fpsyt.2013.00010.

Agcaoglu, O., Wilson, T. W., Wang, Y., Stephen, J., and Calhoun, V. D. (2019). Resting state connectivity differences in eyes open versus eyes closed conditions. Human brain mapping 40, 2488–2498.

Alexopoulos, G. S., Hoptman, M. J., Kanellopoulos, D., Murphy, C. F., Lim, K. O., and Gunning, F. M. (2012). Functional connectivity in the cognitive control network and the default mode network in late-life depression. Journal of affective disorders 139, 56–65.

Alexopoulos, G. S., Hoptman, M. J., Yuen, G., Kanellopoulos, D., Seirup, J. K., Lim, K. O., et al. (2013). Functional connectivity in apathy of late-life depression: a preliminary study. Journal of affective disorders 149, 398–405.

Allen, E. A., Damaraju, E., Eichele, T., Wu, L., and Calhoun, V. D. (2018). EEG Signatures of Dynamic Functional Network Connectivity States. Brain Topography 31, 101–116. doi:10.1007/s10548-017-0546-2.

Allen, E. A., Damaraju, E., Plis, S. M., Erhardt, E. B., Eichele, T., and Calhoun, V. D. (2014). Tracking whole-brain connectivity dynamics in the resting state. Cerebral Cortex 24, 663–676. doi:10.1093/cercor/bhs352.

Bai, T., Wei, Q., Xie, W., Wang, A., Wang, J., Gong-Jun, J. I., et al. (2019). Hippocampal-subregion functional alterations associated with antidepressant effects and cognitive impairments of electroconvulsive therapy. Psychological medicine 49, 1357–1364.

Bai, Y.-M., Chen, M.-H., Hsu, J.-W., Huang, K.-L., Tu, P.-C., Chang, W.-C., et al. (2020). A comparison study of metabolic profiles, immunity, and brain gray matter volumes between patients with bipolar disorder and depressive disorder. Journal of neuroinflammation 17, 42.

Chen, N., Shi, J., Li, Y., Ji, S., Zou, Y., Yang, L., et al. (2021). Decreased dynamism of overlapping brain sub-networks in Major Depressive Disorder. Journal of psychiatric research 133, 197–204.

Clasen, P. C., Beevers, C. G., Mumford, J. A., and Schnyer, D. M. (2014). Cognitive control network connectivity in adolescent women with and without a parental history of depression. Developmental cognitive neuroscience 7, 13–22.

Damaraju, E., Tagliazucchi, E., Laufs, H., and Calhoun, V. D. (2020). Connectivity dynamics from wakefulness to sleep. NeuroImage 220, 117047. doi:10.1016/j.neuroimage.2020.117047.

de Lacy, N., Doherty, D., King, B. H., Rachakonda, S., and Calhoun, V. D. (2017). Disruption to control network function correlates with altered dynamic connectivity in the wider autism spectrum. NeuroImage: Clinical 15, 513–524. doi:10.1016/j.nicl.2017.05.024.

Du, Y., Fu, Z., Sui, J., Gao, S., Xing, Y., Lin, D., et al. (2020). NeuroMark: An automated and adaptive ICA based pipeline to identify reproducible fMRI markers of brain disorders. NeuroImage: Clinical 28, 102375. doi:10.1016/j.nicl.2020.102375.

Ebneabbasi, A., Mahdipour, M., Nejati, V., Li, M., Liebe, T., Colic, L., et al. (2021). Emotion processing and regulation in major depressive disorder: A 7T resting‐state fMRI study. Human Brain Mapping 42, 797–810.

Elaheh Zendehrouh, Sendi, M. S. E., Sui, J., Fu, Z., Zhi, D., and Lv, L. (2020). Aberrant Functional Network Connectivity Transition Probability in Major Depressive Disorder. in 42nd Annual International Conference of the IEEE Engineering in Medicine & Biology Society (EMBC), 1493–1496.

Enneking, V., Dzvonyar, F., Dück, K., Dohm, K., Meinert, S., Lemke, H., et al. (2020). Brain Stimulation Brain functional effects of electroconvulsive therapy during emotional processing in major depressive disorder. i. doi:10.1016/j.brs.2020.03.018.

Fu, Z., Caprihan, A., Chen, J., Du, Y., Adair, J. C., Sui, J., et al. (2019). Altered static and dynamic functional network connectivity in Alzheimer’s disease and subcortical ischemic vascular disease: shared and specific brain connectivity abnormalities. Human Brain Mapping 40, 3203–3221. doi:10.1002/hbm.24591.

Gryglewski, G., Baldinger-Melich, P., Seiger, R., Godbersen, G. M., Michenthaler, P., Klöbl, M., et al. (2019). Structural changes in amygdala nuclei, hippocampal subfields and cortical thickness following electroconvulsive therapy in treatment-resistant depression: longitudinal analysis. The British Journal of Psychiatry 214, 159–167.

Heijnen, W. T., Birkenhäger, T. K., Wierdsma, A. I., and van den Broek, W. W. (2010). Antidepressant pharmacotherapy failure and response to subsequent electroconvulsive therapy: a meta-analysis. Journal of clinical psychopharmacology 30, 616–619.

Jiao, K., Xu, H., Teng, C., Song, X., Xiao, C., Fox, P. T., et al. (2020). Connectivity patterns of cognitive control network in first episode medication-naive depression and remitted depression. Behavioural brain research 379, 112381.

Kennis, M., Gerritsen, L., van Dalen, M., Williams, A., Cuijpers, P., and Bockting, C. (2020). Prospective biomarkers of major depressive disorder: a systematic review and meta-analysis. Molecular psychiatry 25, 321–338.

Leaver, A. M., Wade, B., Vasavada, M., Hellemann, G., Joshi, S. H., Espinoza, R., et al. (2018). Fronto-temporal connectivity predicts ECT outcome in major depression. Frontiers in psychiatry 9, 92.

Leaver, A., Vasavada, M., Sahib, A. K., Kubicki, A., Joshi, S., Woods, R. P., et al. (2020). Modulation of amygdala reactivity following rapidly acting interventions for major depression. 1–12. doi:10.1002/hbm.24895.

Li, G., Liu, Y., Zheng, Y., Li, D., Liang, X., Chen, Y., et al. (2020). Large‐scale dynamic causal modeling of major depressive disorder based on resting‐state functional magnetic resonance imaging. Human brain mapping 41, 865–881.

Liu, G., Jiao, K., Zhong, Y., Hao, Z., Wang, C., and Xu, H. (2021). The alteration of cognitive function networks in remitted patients with major depressive disorder: an independent component analysis. Behavioural Brain Research 400, 113018. doi:10.1016/j.bbr.2020.113018.

Liu, Y., Chen, Y., Liang, X., Li, D., Zheng, Y., Zhang, H., et al. (2020). Altered Resting-State Functional Connectivity of Multiple Networks and Disrupted Correlation With Executive Function in Major Depressive Disorder. Frontiers in Neurology 11, 1–11. doi:10.3389/fneur.2020.00272.

Luo, L., Wu, H., Xu, J., Chen, F., Wu, F., Wang, C., et al. (2021). Abnormal large-scale resting-state functional networks in drug-free major depressive disorder. Brain Imaging and Behavior 15, 96–106. doi:10.1007/s11682-019-00236-y.

Menon, V. (2015). Large-scale functional brain organization. Brain mapping, 449–459.

Mo, Y., Wei, Q., Bai, T., Zhang, T., Lv, H., and Zhang, L. (2020). Bifrontal electroconvulsive therapy changed regional homogeneity and functional connectivity of left angular gyrus in major depressive disorder. Psychiatry Research 294, 113461. doi:10.1016/j.psychres.2020.113461.

Mulders, P. C. R., van Eijndhoven, P. F. P., Pluijmen, J., Schene, A. H., Tendolkar, I., and Beckmann, C. F. (2016). Default mode network coherence in treatment-resistant major depressive disorder during electroconvulsive therapy. Journal of affective disorders 205, 130–137.

Mulders, P. C., van Eijndhoven, P. F., Schene, A. H., Beckmann, C. F., and Tendolkar, I. (2015). Resting-state functional connectivity in major depressive disorder: A review. Neuroscience and Biobehavioral Reviews 56, 330–344. doi:10.1016/j.neubiorev.2015.07.014.

Organization, W. H. (2008). The global burden of disease: 2004 update. World Health Organization.

Ozomaro, U., Wahlestedt, C., and Nemeroff, C. B. (2013). Personalized medicine in psychiatry: problems and promises. BMC medicine 11, 1–35.

Perlman, K., Benrimoh, D., Israel, S., Rollins, C., Brown, E., Tunteng, J.-F., et al. (2019). A systematic meta-review of predictors of antidepressant treatment outcome in major depressive disorder. Journal of affective disorders 243, 503–515.

Perrin, J. S., Merz, S., Bennett, D. M., Currie, J., Steele, D. J., Reid, I. C., et al. (2012). Electroconvulsive therapy reduces frontal cortical connectivity in severe depressive disorder. Proceedings of the National Academy of Sciences 109, 5464–5468.

Posner, J., Cha, J., Wang, Z., Talati, A., Warner, V., Gerber, A., et al. (2016). Increased Default Mode Network Connectivity in Individuals at High Familial Risk for Depression. Neuropsychopharmacology 41, 1759–1767. doi:10.1038/npp.2015.342.

Qiu, H., Li, X., Luo, Q., Li, Y., Zhou, X., Cao, H., et al. (2019). Alterations in patients with major depressive disorder before and after electroconvulsive therapy measured by fractional amplitude of low-frequency fluctuations (fALFF). Journal of affective disorders 244, 92–99.

Repple, J., Mauritz, M., Meinert, S., de Lange, S. C., Grotegerd, D., Opel, N., et al. (2020). Severity of current depression and remission status are associated with structural connectome alterations in major depressive disorder. Molecular psychiatry 25, 1550–1558.

Rush, A. J., Trivedi, M. H., Wisniewski, S. R., Nierenberg, A. A., Stewart, J. W., Warden, D., et al. (2006). Acute and longer-term outcomes in depressed outpatients requiring one or several treatment steps: a STAR* D report. American Journal of Psychiatry 163, 1905–1917.

Sartorius, A., Demirakca, T., Böhringer, A., von Hohenberg, C. C., Aksay, S. S., Bumb, J. M., et al. (2019). Electroconvulsive therapy induced gray matter increase is not necessarily correlated with clinical data in depressed patients. Brain stimulation 12, 335–343.

Schlösser, R. G. M., Wagner, G., Koch, K., Dahnke, R., Reichenbach, J. R., and Sauer, H. (2008). Fronto-cingulate effective connectivity in major depression: a study with fMRI and dynamic causal modeling. Neuroimage 43, 645–655.

Sendi, M. S. E., Zendehrouh, E., Ellis, C. A., Liang, Z., Fu, Z., Mathalon, D. H., et al. (2021a). Aberrant Dynamic Functional Connectivity of Default Mode Network in Schizophrenia and Links to Symptom Severity. Front. Neural Circuits 15.

Sendi, M. S. E., Zendehrouh, E., Miller, R. L., Fu, Z., Du, Y., Liu, J., et al. (2021b). Alzheimer’s Disease Projection From Normal to Mild Dementia Reflected in Functional Network Connectivity: A Longitudinal Study. Front. Neural Circuits 14. doi:10.3389/fncir.2020.593263.

Sendi, M. S. E., Zendehrouh, E., Sui, J., Fu, Z., Zhi, D., Lv, L., et al. (2021c). Aberrant dynamic functional connectivity of default mode network predicts symptom severity in major depressive disorder. Brain Connectivity, 1–36. doi:10.1089/brain.2020.0748.

Settell, M. L., Testini, P., Cho, S., Lee, J. H., Blaha, C. D., Jo, H. J., et al. (2017). Functional circuitry effect of ventral tegmental area deep brain stimulation: imaging and neurochemical evidence of mesocortical and mesolimbic pathway modulation. Frontiers in neuroscience 11, 104.

Sharp, R. (2015). The hamilton rating scale for depression. Occupational Medicine 65, 340. doi:10.1093/occmed/kqv043.

Sheline, Y. I., Price, J. L., Yan, Z., and Mintun, M. A. (2010). Resting-state functional MRI in depression unmasks increased connectivity between networks via the dorsal nexus. Proceedings of the National Academy of Sciences 107, 11020–11025.

Stange, J. P., Bessette, K. L., Jenkins, L. M., Peters, A. T., Feldhaus, C., Crane, N. A., et al. (2017). Attenuated intrinsic connectivity within cognitive control network among individuals with remitted depression: temporal stability and association with negative cognitive styles. Human brain mapping 38, 2939–2954.

Takamiya, A., Chung, J. K., Liang, K., Graff-Guerrero, A., Mimura, M., and Kishimoto, T. (2018). Effect of electroconvulsive therapy on hippocampal and amygdala volumes: systematic review and meta-analysis. The British Journal of Psychiatry 212, 19–26.

Tsuchiyagaito, A., Smith, J. L., El-sabbagh, N., Zotev, V., Misaki, M., Al, O., et al. (2021). NeuroImage: Clinical Real-time fMRI neurofeedback amygdala training may influence kynurenine pathway metabolism in major depressive disorder. NeuroImage: Clinical 29, 102559. doi:10.1016/j.nicl.2021.102559.

Van Waarde, J. A., Scholte, H. S., Van Oudheusden, L. J. B., Verwey, B., Denys, D., and Van Wingen, G. A. (2015). A functional MRI marker may predict the outcome of electroconvulsive therapy in severe and treatment-resistant depression. Molecular psychiatry 20, 609–614.

Vasic, N., Walter, H., Sambataro, F., and Wolf, R. C. (2009). Aberrant functional connectivity of dorsolateral prefrontal and cingulate networks in patients with major depression during working memory processing. Psychological medicine 39, 977.

Veer, I. M., Beckmann, C., Van Tol, M.-J., Ferrarini, L., Milles, J., Veltman, D., et al. (2010). Whole brain resting-state analysis reveals decreased functional connectivity in major depression. Frontiers in systems neuroscience 4, 41.

Wang, H., Li, R., Zhou, Y., Wang, Y., Cui, J., Nguchu, B. A., et al. (2018a). Altered cerebro-cerebellum resting-state functional connectivity in HIV-infected male patients. Journal of neurovirology 24, 587–596.

Wang, J., Wang, Y., Huang, H., Jia, Y., Zheng, S., Zhong, S., et al. (2020). Abnormal dynamic functional network connectivity in unmedicated bipolar and major depressive disorders based on the triple-network model. Psychological medicine 50, 465–474.

Wang, J., Wei, Q., Bai, T., Zhou, X., Sun, H., Becker, B., et al. (2017). Electroconvulsive therapy selectively enhanced feedforward connectivity from fusiform face area to amygdala in major depressive disorder. Social cognitive and affective neuroscience 12, 1983–1992.

Wang, J., Wei, Q., Wang, L., Zhang, H., Bai, T., Cheng, L., et al. (2018b). Functional reorganization of intra‐and internetwork connectivity in major depressive disorder after electroconvulsive therapy. Human Brain Mapping 39, 1403–1411.

Wei, Q., Bai, T., Chen, Y., Ji, G., Hu, X., Xie, W., et al. (2018). The changes of functional connectivity strength in electroconvulsive therapy for depression: a longitudinal study. Frontiers in neuroscience 12, 661.

Williams, L. M., Coman, J. T., Stetz, P. C., Walker, N. C., Kozel, F. A., George, M. S., et al. (2021). Identifying response and predictive biomarkers for Transcranial magnetic stimulation outcomes: protocol and rationale for a mechanistic study of functional neuroimaging and behavioral biomarkers in veterans with Pharmacoresistant depression. BMC psychiatry 21, 1–17.

Yan, C. G., Chen, X., Li, L., Castellanos, F. X., Bai, T. J., Bo, Q. J., et al. (2019). Reduced default mode network functional connectivity in patients with recurrent major depressive disorder. Proceedings of the National Academy of Sciences of the United States of America 116, 9078–9083. doi:10.1073/pnas.1900390116.

Yang, Z., Guo, H., Ji, S., Li, S., Fu, Y., Guo, M., et al. (2020). Reduced Dynamics in Multivariate Regression-based Dynamic Connectivity of Depressive Disorder. in 2020 IEEE International Conference on Bioinformatics and Biomedicine (BIBM) (IEEE), 1197–1201.

Yoav Benjamini ; Yosef Hochberg (1995). Controlling the False Discovery Rate: A Practical and Powerful Approach to Multiple Testing. Royal Statistical Society. Series B (Methodological) 57, 289–300.

